# Trait evolution under multiple selection pressures: Prey responses to predictable and unpredictable variation

**DOI:** 10.1101/816314

**Authors:** Manvi Sharma, Kavita Isvaran

## Abstract

When a strong selection pressure, such as predation risk, varies widely in space and time, how should prey respond? When risk varies predictably, prey are hypothesized to respond in a risk-sensitive manner. It is less clear how prey should respond when risk varies unpredictably.

Additionally, prey response may also depend on how predation risk interacts with other selection pressures. Our understanding of the complex action of multiple and variable selection pressures on prey traits is still comparatively poor. Here, we examine how predictable and unpredictable aspects of predation risk act together with another important selection pressure to influence prey behaviour in the rock pool breeding mosquito, *Aedes vexans.* Through the selection of sites for oviposition, female mosquitoes can influence the predation risk faced by their offspring. We tested how females select oviposition sites, when encountering pools that vary in larval predation risk and desiccation risk. We comprehensively quantified spatial and temporal variation in predation risk by measuring densities of predatory dragonfly nymphs in rock pools of different sizes, along the mosquito breeding season. We also measured hydroperiod length. We next conducted manipulative experiments in rock pools and measured female oviposition responses to variation in predation and desiccation risks. Predation risk varied widely in space and time. Desiccation risk only appeared important for the small pools. Ovipositing females appeared to respond to these multiple aspects of variation in selection pressures. Females seemed to respond to predictable variation by avoiding large pools that permanently harboured predators in natural settings. Female responses were more variable to medium-sized pools with naturally stochastic predator densities, highlighting the role of unpredictability in predation risk in shaping behaviour. Females did not clearly prefer small pools that were naturally devoid of predators but carried high desiccation risk, suggesting that they balance multiple risks – predation versus desiccation – when choosing oviposition sites. Our study suggests that wild populations may commonly experience multiple and variable selection pressures that can favour seemingly puzzling trait variation. We highlight the need to quantify variation in selection pressures and investigate how such variation, especially the unpredictable aspects, shapes prey traits.

## INTRODUCTION

In wild populations, traits are likely to evolve in response to multiple selection pressures, which may act complementarily or in opposing directions with different strengths (Andersson, 1982; Templeton & Shriner, 2004; Heinen-kay et al., 2015; Labarbera, Hayes, Marsh, & Lacey, 2017). We have evidence from several studies that not only metric traits, such as body colour (Endler, 1983; Heinen-kay et al., 2015) and body size (Anholt, 1991), but also behavioural traits, such as, courtship displays (Endler, 1987; Sih 1994) and activity levels (Brodin & Johansson, 2004), are likely to be shaped by multiple selection pressures. Of the different selection pressures influencing a trait, certain selection pressures, such as risk of predation, can impose a stronger force than other on the process of trait evolution (Endler, 1987). When a strong selection pressure, such as predation risk, varies widely in space and time, how should prey animals tailor response towards this variation?

The solution to the problem is likely to depend on whether the variation in predation risk in the environment, is *predictable*, or not. Theory and empirical work suggests that when the intensity of predation risk in the environment varies *predictably*, prey should evolve to detect and track the variation in predation risk, and respond in a risk-sensitive manner (Helfman, 1989; Roux, Diabate, & Simard, 2014). Prey can gain information on predation risk directly, through predator odour and predator encounter (Cronin, 2004; Laurila, Järvi-Laturi, Pakkasmaa, & Merilä, 2004; Ferrari, Wisenden, & Chivers, 2010; Gulsby, Cherry, Johnson, Conner, & Miller, 2018), or indirectly, through environmental characteristics that predict predation risk, such as refuge site availability or lunar intensity (reviewed in (Verdolin, 2006; Abramsky, Rosenzweig, Belmaker, & Bar, 2004). Much less is known about how prey should respond to *unpredictable* variation, that is where prey are unable to detect and track variation in predation risk, or to environments with both predictable and unpredictable sources of variation in predation risk (Sih, 1992; Bouskila, Blumstein, & Mangel, 1995; Martin, 2011).

Unpredictable variation in predation risk has been studied previously only in the form of prey responses to foreign/novel predators. These studies tested immediate prey responses to novel predators (Brown, Ferrari, Elvidge, Ramnarine, & Chivers, 2013; Ferrari et al., 2015) on an ecological time scale. Surprisingly, no empirical studies have tested how prey responses should evolve when prey is exposed to unpredictably varying predation risk at an evolutionary time scale. Studies on behaviour, in contexts other than predation risk, such as mating strategies and nesting time decisions, suggest that unpredictable variation in selection factors could lead to the evolution of alternative strategies, such as bet-hedging (Philippi & Seger, 1989; Simons, 2011; Starrfelt & Kokko, 2012). To our knowledge, unpredictable variation in the context of predation, has not received much attention, beyond theoretical models (Luttbeg & Sih, 2010; Feyten & Brown, 2018).

To study prey response to both predictable and unpredictable variation in predation risk in the wild, we need to first measure this natural variation in risk of predation over a period of time.This can be challenging as predators are typically present at low densities (Boland, 2003; Creel & Winnie, 2005), but see (Juliana, Kotler, Brown, Mukherjee, & Bouskila, 1999). However, recent evidence highlights the importance of measuring this variation, because they suggest that the risk of predation changes dramatically in the wild over space and time, for example, when predators move across a landscape (Thaker et al., 2011; Watts, Jones, Herrig, Miller, & Tenhumberg, 2018), or when predators reproduce (Yoshida, Jones, Ellner, Fussmann, & Hairston, 2003; Deacon, Jones, & Magurran, 2018). Studies on the ecology of fear and predator-prey space use, measure and report systematic changes in predation risk to study prey behavioural response (Hammond, Luttbeg, & Sih, 2007; Mukherjee, Zelcer, & Kotler, 2009), (Thaker et al., 2011). These studies typically assume that prey can reliably detect changes in predation risk. There is an exciting unexplored scenario where predation risk varies stochastically and prey cannot, therefore, reliably detect changes in predation. In our study, we overcome the challenges of quantifying variation in predation risk in the wild by using a predator-prey model system that is tractable in the wild. We use *Aedes vexans, a* mosquito that breeds in ephemeral pools, and *Bradinopygea geminata*, a dragonfly, as the prey-predator model system. We focus on the anti-predator behaviour shown by female mosquitoes towards predation risk faced by their offspring from predatory dragonfly nymphs. We quantify and then test how predictable and unpredictable variation in predation risk from *B. geminata* influences how *Ae. vexans* females select sites for oviposition

Patch-breeding animals, such as frogs and mosquitoes, typically encounter pools that vary in multiple risk factors. The risk of offspring predation (Rieger, Binckley, & Resetarits Jr., 2004; Silberbush & Blaustein, 2011) is thought to be an important selection pressure that influences the selection by females of suitable sites for ovipositing. *Aedes vexans*, which breeds in ephemeral pools (Balasubramanian & Balasubramanian, 2013), such as rock-pools, is also reported to respond to larval predation risk. A meta-analysis of mosquito oviposition response to larval predators showed that *Aedes* spp., unlike other mosquito species, showed substantial variation in their response (Vonesh & Blaustein, 2010). Interestingly, the oviposition behaviour of *Aedes* spp. may contribute towards unpredictability in larval predation risk. Females deposit eggs along the moist edges of pools, above the surface of water and these eggs hatch only when inundated by rain-water following an event of rainfall (Clements. 1990). Consequently, there is an element of uncertainty as the level of predation risk encountered by females at the time of oviposition might not represent the level of risk that her offspring experience when her eggs eventually hatch. Apart from predation risk, the offspring of pool breeding animals are particularly susceptible to risk of mortality due to pool desiccation (Brady & Griffiths, 2000) and thus, pool hydroperiod length is also likely to influence female oviposition behaviour of *Aedes* spp.

In our study, using *Ae. vexans* and *Bradinopygea*, as the prey-predator model system, in a natural field setting, we investigated how oviposition site selection is influenced by different aspects of variation in larval predation risk. We also incorporated the influence of pool desiccation risk on oviposition site selction decisions. Specifically, we asked 1) What is the nature of variation (predictable and unpredictable) in larval predation risk that ovipositing females face along the mosquito breeding season, i.e., identify pools that reliably or unreliably predicted predator presence, 2) How do ovipositing females respond to predictable and unpredictable variation in predation risk in a natural field setting. 3) How do females respond to risk of pool desiccation when selecting sites for oviposition? To address these questions, we first measured variation in larval predation risk (i) over space (i.e., among different pools) and (ii) over time (i.e., at different times during the mosquito breeding season). We hypothesised that females would be sensitive to both predictable and unpredictable variation in predation risk. We expected females to respond to predictable variation in predation risk by ovipositing in a risk-sensitive manner - 1) females should respond to spatial variation in predation risk by showing stronger avoidance of pools with higher predation risk; 2) females should respond to systematic temporal variation in risk by showing a stronger avoidance during the period in the mosquito breeding period when predation risk is highest. Additionally, we expected females to respond to unpredictable variation in predation risk and predicted a lack of a consistent avoidance of pools associated with unpredictability in predation risk. Along with larval predation risk, ovipositing females in the wild face risk of pool desiccation. Together with larval predation risk, we also quantified desiccation risk of oviposition sites and tested female response to pools varying in desiccation risk. We expected that pools that could potentially dry up before the completion of mosquito larval maturation and therefore, present a high risk of larval mortality, would be avoided by ovipositing females throughout the mosquito breeding period.

## METHODS

### Characterising predictable and unpredictable variation in predation risk

We carried out all work at the Rishi valley school campus (13.63, 78.45), located in Madanapalle, Andhra Pradesh, India. Our study site primarily has rocky outcrops with thorny scrub vegetation. The field site receives most of the rainfall between June to December, and September has the highest probability of rain (Supplementary Fig. S1)The school campus has several sheet rocks with depressions that get inundated with rain water forming ephemeral rock pools. A preliminary survey of rock pool biodiversity sampling at the field site revealed that *Aedes vexans* was the dominant rock pool breeding mosquito species and nymphs of the granite ghost dragonfly, *Bradinopyga geminata* are the dominant aquatic predators. *Ae. vexans*, is a typical floodwater mosquito. It oviposits eggs in temporary pools and the eggs can undergo dormancy and hatch after a flooding event (Clements 1990). *Ae. vexans* is a day-biting mosquito and is known to be a vector of Tahyna virus in Central Europe (Pilaski, 1987) and Rift Valley Fever virus in West Africa (Zeller, Fontenille, Traore-Lamizana, Thiongane, & Digoutte, 1997). Females are reported to live for 3 to 6 weeks (Horsfall et al. 1973) in lab conditions; females typically have a body size at emergence of 3 −4.45 mm; and carry much less protein and lipid reserves at emergence, compared to other related mosquitoes, such as *Ae. aegypti* (Briegel, 1990; Briegel, Waltert, & Kuhn, 2001). The number of mature eggs in wild *Ae. vexans* females varies between 108 to 180, with an average of 132 (Horsfall et al. 1973). The rock pools get populated with diverse rock pool fauna, such as Hydrophilidae (water beetles), Chironomidae larvae (midge larvae), Nepidae (water scorpions), Copepods, Culicidae (mosquito larvae), Notonectidae (backswimmers), Odonate larvae (dragonfly larvae) and Dichroglossidae (skittering frogs) and Bufonidae tadpoles.

To estimate patterns of variation in the risk of predation, we sampled all pools (N=89), that were located on the largest sheet rock, repeatedly during the mosquito breeding season, from July to December 2015. To estimate predator density in a pool, we used a sampling grid of 50 x 50 cm area and placed it over the pool, scooped out the nymphs of *Bradinopyga geminata*, falling within the sampling area into a container using a fishing net, and counted the removed nymphs. During a sampling occasion, we also recorded presence/absence data for other rock pool fauna that were collected in the fishing net: tadpoles, copepods, mosquito larvae, backswimmers and water beetles. Each pool was sampled once a month from July to December, yielding a dataset of 534 samples. A few pools were found to be filled with cow dung and were excluded from the analyses.

### Characterising desiccation risk in rock-pools

Rock pools of variable size were expected to be associated with different levels of desiccation risk as their water holding capacity would vary with size. For patch breeding organisms, hydroperiod length, the number of days for which a pool can hold water before it dries, is used as a measure of desiccation risk (Resetarits, 1996; Brady & Griffiths, 2000). We calculated hydroperiod length as the number of days from the last heavy shower (>5mm), until the pool dried. The pools were monitored every day following a heavy shower and dry/not dry status was recorded. The session was terminated either when all the pools under observation dried out or when an event of rainfall occurred; in the latter case, a new session was then started. The duration of a session ranged from 3 days (when rain occurred 3 days after the session was started) to a 17 day long uninterrupted sampling session. These observations were recorded in 2015 following an event of rainfall on 6 occasions, from August to December. The dataset consisted of 89 pools sampled 6 times representing a sample size of 534 data points. The few pools that were found to be filled with cow dung were excluded from the analyses.

### Measuring adult oviposition response

To test how ovipositing females in the wild respond to patterns of predictable and unpredictable variation in predation risk, and if they pay attention to desiccation risk, we carried out a field experiment. For a sub-set of pools from those characterised above, we experimentally manipulated predation risk by adding and removing predators. We then measured oviposition responses to these experimental pools. These experimental pools were a sub-set of the larger background level of spatial and temporal variation in predation risk and desiccation risk in the environment. We cleaned pools using a fishing net prior to adding treatments for the manipulation experiments. Post cleaning, we covered the pools with a mesh of appropriate size that curtailed oviposition by adult dragonflies but permitted oviposition by adult mosquitoes. The pool water was allowed to stand for 2 days after which pools where either assigned to a predator treatment or to a control treatment. In the predator treatment pools, predators (*Bradinopyga* nymphs) were added. The number of predators added to a pool was chosen to represent average predator densities that were found to be naturally occurring for a given pool size from a survey done in June 2014. The predators were collected from nearby pools that were not a part of the experiment. In the course of a trial, predators found dead in the pool were immediately replaced. For control pools, no predators were added. We tested a total of 31 unique pools ranging from 20 to 320 cm perimeter. Pools were chosen covering a range of sizes to represent the natural range of pool sizes that *Aedes* spp. oviposits in. This range of pool sizes also captured the natural variation in larval predation risk (described in detail in the Results section).

To quantify female oviposition preferences, we collected and counted the eggs laid by females in experimental pools. *Aedes* females oviposit along the walls of the rock pools (Clements, 1999) above the water line. We calculated the perimeter of rock pools using photographs and image manipulation software ImageJ (Schneider et. al. 2012). For collecting eggs, we placed sheets of white cloth (ovistrips) of width 7.5 cm and of length equal to 50% of perimeter of the pool to be sampled (with the aim to sample pools in proportion to their size). The region of the pool perimeter where the strips were placed was randomised between trials. We collected ovistrips after 24 hours and attached fresh ovistrips along the wall of the pool each day. This 24 hour period constituted a trial. These trials were conducted over a period of 15 days called a session. After termination of a session, the predator/control treatments were switched so that the same pool received predator as well as control treatment. This exercise would account for pool identity related biases in the experimental design. The collected ovistrips were dried at the field station house and eggs were counted using a microscope. A subset of eggs from each trial was soaked for hatching and the larvae and adults were used for species identification. We used morphological keys from Walter Reed Biosystems and Florida entomological guide for species identification (http://www.wrbu.org/VecID_MQ.html, http://fmel.ifas.ufl.edu/fmel---mosquito-key/). Trials were run from July to December 2014 and 2015 which covered two mosquito breeding seasons. A total of 95 trials was carried out during the study period, out of which 70 were successful, that is, the trials that were not interrupted by heavy rainfall.

### Statistical analysis

#### Predictable and unpredictable variation in larval predation risk

To examine patterns in variation in predation risk over time and space (pools varying in size), we ran a generalised linear mixed effects model with negative binomial errors with zero-inflation assumed to be constant. We modelled predator density as the response variable and predictor variables chosen were log of pool perimeter, month, and the interaction term between month and log of pool perimeter. Pool identity was included as a random effect in the model. The interaction term was included to model the combined effect of month and pool size on predator density, that could potentially arise because of patterns in predator breeding behaviour. We did these analyses using the R package glmmTMB (Brooks et.al. 2019). In addition to formally modelling spatial and temporal variation in predator density, a quantitative measure, we calculated the probability of predator presence as a function of pool size to test how reliably pool size could predict predator presence in the mosquito breeding season. This measure was calculated because of the unique life history of *Ae. vexans* - eggs hatch only following a rainfall event (Clements, 1990). Hence, for ovipositing females, the current predator density of the pool might not be a reliable measure of risk of predation to her offspring. Thus we calculated probability of predator presence for pools of different size classes. We collapsed the pool perimeter into 10 equal size classes (L1(<50cm), L2(50-100cm), L3(100-150cm), L4(150-200cm), L5(200-250cm), L6(250-300cm), L7(300-350cm), L8(350-400cm), L9(400-450cm), and L10(>450cm). We calculated the probability of predator presence as the number of pools in a given size class with at least one predator recorded, divided by the total number of pools. For calculating probability of predator presence, we pooled together the samples from all six months (i.e. n = 427 predator presence/absence observations from 89 pools). 95% bootstrapped confidence intervals were calculated for the probability of predator presence for each size class by resampling pools within each size class.

### Variation in pool desiccation risk

For modelling pool desiccation risk, we recorded hydroperiod length (number of days for the pool to dry up after a rainfall event). To estimate pool desiccation risk from hydroperiod length data we used a survivor analysis approach and modelled desiccation risk as the probability of drying out. This approach allowed the use of censored data, i.e, in this case, pools whose dry/not dry status is known for a certain time period even though the final outcome, i.e., hydroperiod length, is not known. This approach was used because we terminated our sampling before the large pools could dry out completely. We ran a mixed effects cox-proportional hazards model with hydroperiod length as a response variable and log pool perimeter, month and interaction term between month and log pool perimeter as explanatory variables. The interaction term was included in the model because microclimatic variables, such as sunlight, cloud cover, relative humidity, and daily temperature could affect how hydroperiod length changes with pool size and month. July and November could not be included in the analyses as hydroperiod length data could not be collected for those months. These analyses were done using the coxme package in R (Therneau, 2015). We checked model assumptions of the cox-proportional hazard model using the zph test in a simpler model without random effects.

### Measuring female oviposition behaviour

We used two indices to measure female oviposition behaviour; 1) Egg density – Number of eggs laid per unit of perimeter sampled in a pool (this assumes that if females show no preference, the number of eggs laid in a pool should be proportional to the perimeter of the pool, and 2) Coefficient of variation (cv) in egg densities – calculated as (standard deviation/mean * 100), can be used as a measure of relative variability between treatments.

We ran a linear mixed effects model (LMM) with normal errors with mean egg density in a session (mean of all trials in that session for that pool) as response variable, and pool size, predator treatment (presence/absence), background predation risk and all two-way interaction terms as explanatory variables. To examine the influence of temporal patterns in predator density, we included a two-factor categorical variable called background predation risk. Based on our findings on patterns of variation in predation risk (see Fig. 1), we selected October and November to represent “high” background predation risk, while the rest of the mosquito breeding period represented “low” background predation risk in the model. We modelled pool size as a categorical rather than a continuous variable because we had no clear expectation for the underlying shape of the relationship between response variable (egg densities) and pool size. As a result, we categorised pool size into 4 size classes to capture spatial variation in predator density that occurs naturally among pools. This was based on our findings on patterns of variation in predation risk (see supplementary table 1)). Due to large differences in variance of egg densities, between large pools (>300cm) and all other pools (<300cm), the large (level 4) pools were removed from formal analyses. Pool identity was modelled as a random effect to account for the repeated measures on pools. We also categorised the mosquito breeding season into the pre-diapause (August to November), and diapause season (December). This season categorisation was done because the oviposition behavior of *Aedes* spp. has been shown to change in the diapause season i.e. towards the end of the rainfall season (Edgerly, McFarland, & Morgan, 1998). We carried out separate analyses for pre-diapause and diapause season, rather than a single analysis, because studies suggest that female oviposition strategy during the diapause season is different from the rest of the mosquito breeding season (Toma, Severini, Di Luca, Bella, & Romi, 2003). The second index we calculated was coefficient of variation - CV=(standard deviation /mean) * 100 - in egg densities, to test how females respond to unpredictable variation in predation risk. We tested if female response was was more variable towards pools that were naturally associated with unpredictability in predation risk. We calculated CV in egg densities for pools under low and high background predation risk and calculated 95% bootstrap confidence intervals around the CV for interpretation. We calculated CVs only for the pre-diapause season that was from July to November.

**Figure 1:**
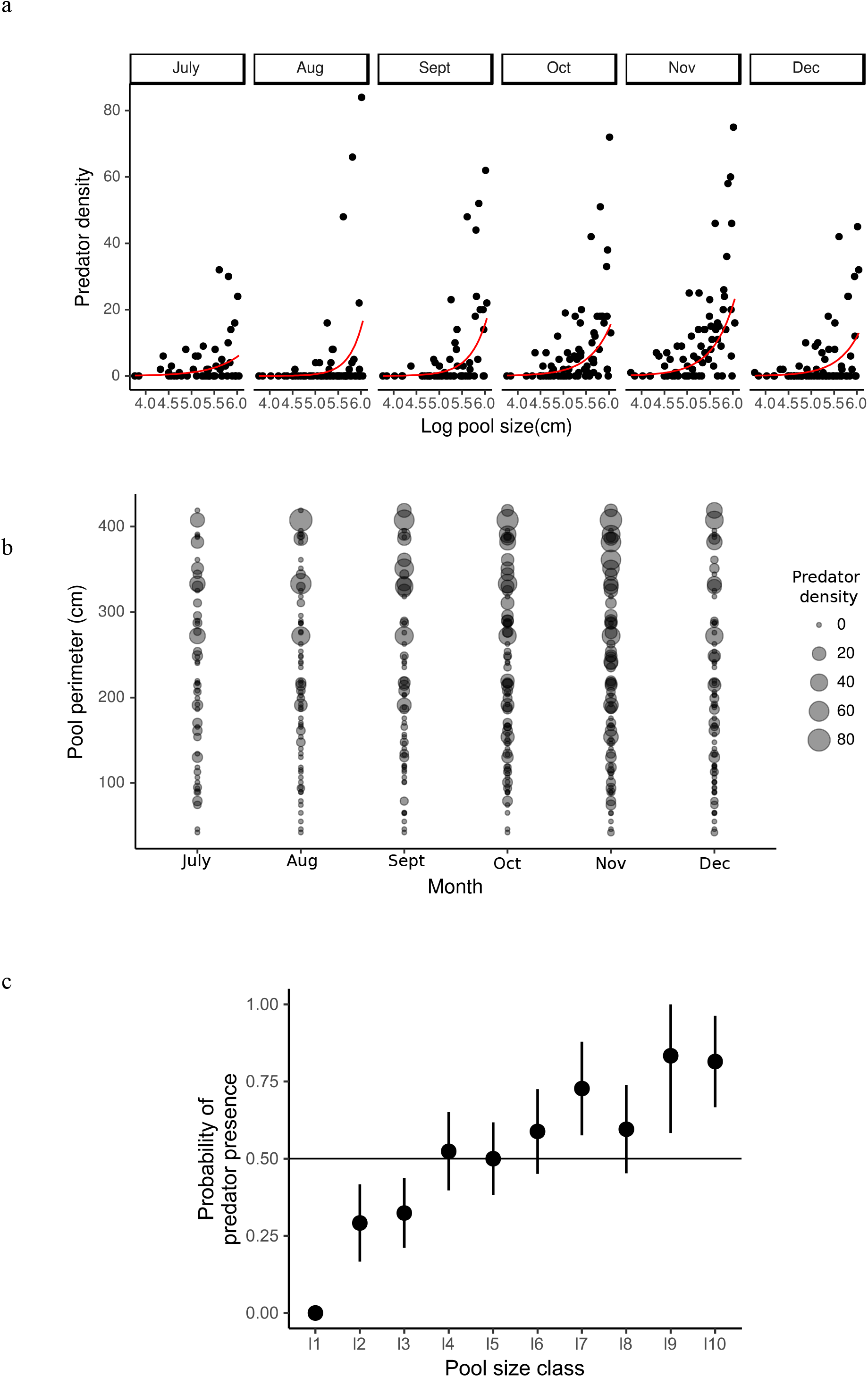
a) Predator density increases with pool size and the slope of this relationship varies among months. Red lines represent predictions from the GLMM (see Table S2). b) Predator density varies in space(along pool size) and in time. Small pools do not harbour predators while large pools harbour high densities of predators throughout the mosquito breeding season from July to December. c) Small and large pools reliably predict predator absence and presence respectively. Medium sized pools (150-250cm) are associated with unpredictable variation in predation risk. Error bars represent means with 95% bootstrapped confidence intervals. The pool size classes based on perimeter measurements were; L1(<50cm), L2(50-100cm), L3(100-150cm), L4(150-200cm), L5(200-250cm), L6(250-300cm), L7(300-350cm), L8(350-400cm), L9(400-450cm), L10(>450cm).

All inferences were based on the full model and we used likelihood ratio tests for testing the statistical significance of fixed effects. Model criticism was performed to check whether model assumptions were satisfied. We carried out all analyses using the statistical software R (v.3.2.2) (R core team 2017). We used the lme4 package in R for running LMMs (Bates et al. 2015). We conducted the female oviposition behaviour experiments from August to December in 2015-2016 resulting in a dataset of n=742 from 28 unique pools. As described earlier, the data were collected during a 10-12 day period called a session, and so we conducted all our analyses using session wise means. Hence, we had a collapsed dataset of n=140.

## RESULTS

### Predictable variation in predation risk

Predator density increased with pool perimeter in all months during the course of the study. The predator density ranged from 0 predators in the smallest pools to 84 predators/2500 cm^2^, in one of the larger pools. The steepness of this positive relationship between pool perimeter and predator density varied between months (GLMM interaction term: *χ^2^* = 13.47, df = 5, *p* <0.05, Table S2) with the predator density increasing with pool size in all months but most steeply in November (Fig. 1).

### Unpredictable variation in predation risk

Probability of predator presence was calculated for 10 equal size classes of pools. Small pools of perimeter < 50cm, had 0 probability of predator presence. On the other hand, large pools with perimeter ranging from 300cm to 450cm, had, on an average, a 60-80% chance of harbouring predators. Interestingly, for many intermediate size classes with perimeter ranging from 150 to 250cm, the probability of predator presence was close to 0.5 (Fig. 1c). Eggs of *Aedes* spp. hatch only when inundated with water following a rainfall event, and thus large pools that consistently harbour predators. Small pools that are consistently devoid of predators, could reliably predict predator presence and absence respectively. Interestingly, medium sized pools on the other hand are stochastic in terms of predator presence as they did not reliably predict predator presence.

### Desiccation risk

There was much variation in hydroperiod length between pools: the smallest pool of perimeter 15cm dried in 2 days (Fig. 2), whereas several large pools did not dry during the course of the sampling session. The results from the mixed effects cox-proportional hazards model suggested that the relative log hazard of drying decreased substantially as pool perimeter increased (Fig. 2). The relative hazard of drying was highest in October and lowest in December (see Table S3). Given that the larvae of *Aedes* spp. mature in 4-6 days (Clements, 1999), we expect only small pools (<50cm) to pose risk of desiccation.

**Figure 2:**
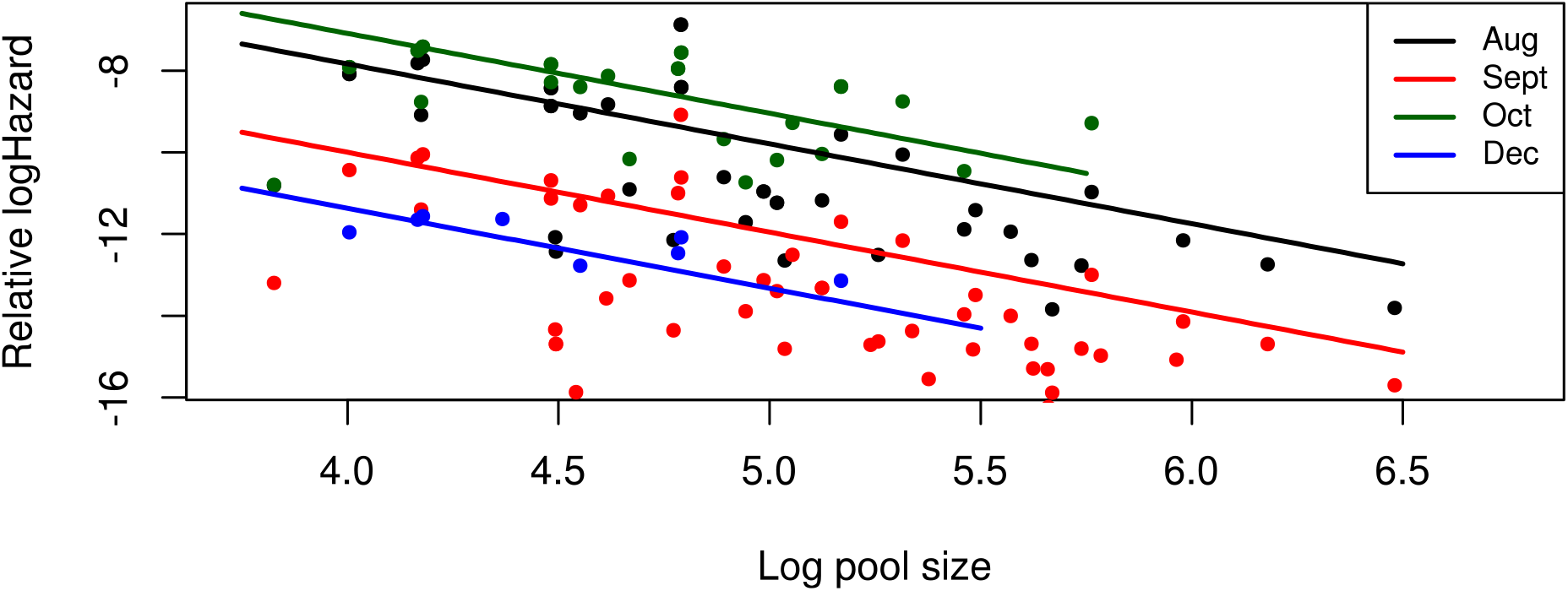
Risk of pool desiccation (Relative log hazard) decreases with pool size and this relationship varies among months. Lines show results from cox-proportional hazards model.

### Female oviposition behaviour

Females appeared to strongly discriminate between pools on the basis of pool size as they avoided pools of large size (>300cm perimeter) throughout the breeding season (Fig. 3a). Due to large differences in variance in egg densities between large pools (>300cm) and all other pools (<300cm), the level 4 pools were removed from formal analyses. LMM results show that in pre-diapause season, the number of eggs laid by females depend on both background predation risk and predator status: when background predation risk is low, females oviposit more eggs in predator-free pools, but when background predation risk is high, this response switches, and females lay more eggs in predator pools (LMM interaction term, *χ*^2^ = 6.77, p<0.01, df=1). In the diapause season, females did not consistently oviposit a higher number of eggs in control pools, compared with predator pools across all pool size classes (LMM interaction term, *χ*^2^ = 0.03, p<0.84, df=1, see Table S4). Overall, the number of eggs laid was higher in diapause season for small and medium pool size classes when compared with pre-diapause season (Fig. 3a).

**Figure 3.**
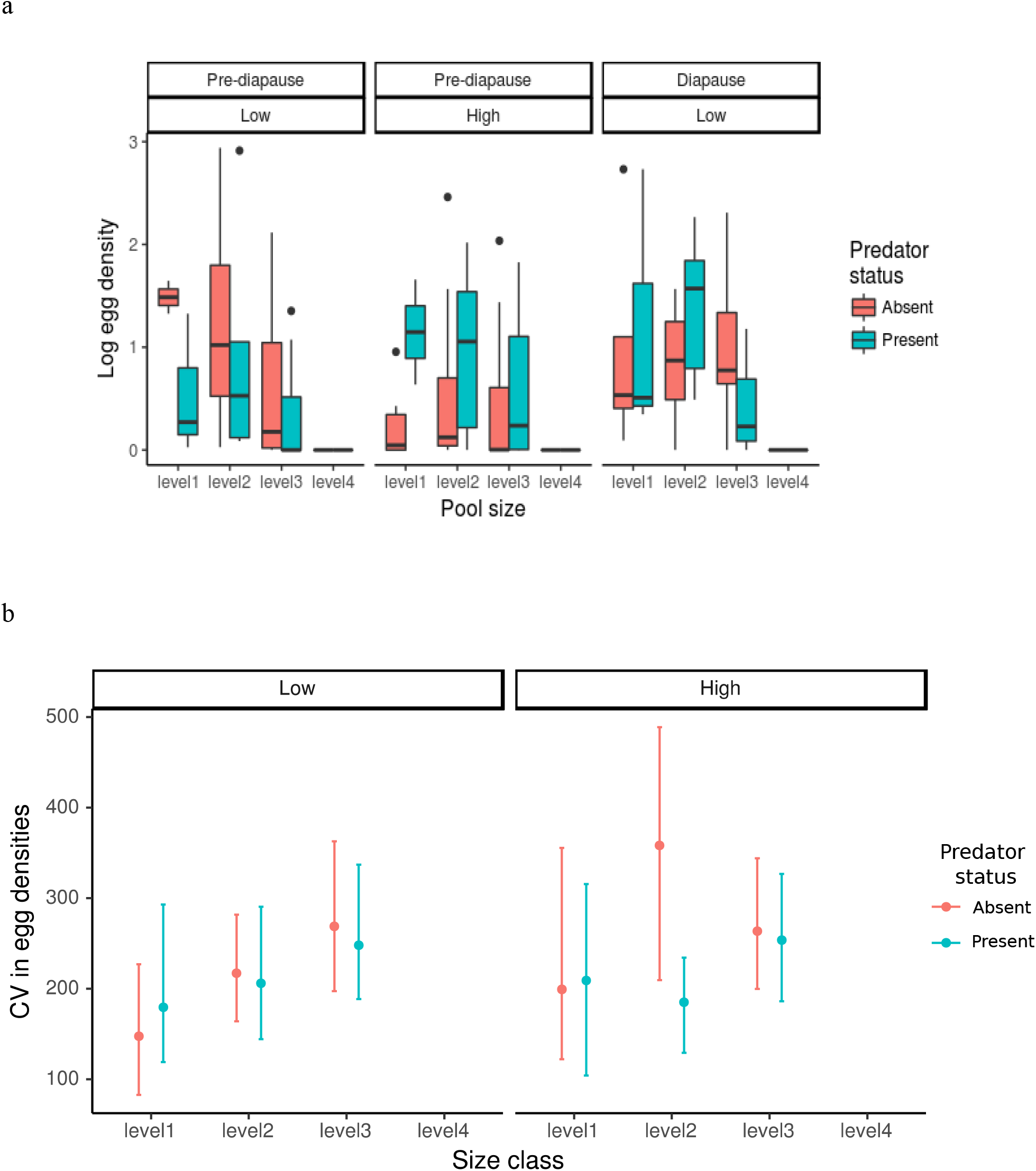
a: Females respond to predictable variation in predation risk; large pools are consistently avoided. Pre-diapause season represented from July to November and Diapause season in December. Low and High represent background predation risk. The different levels indicate pool size classes; level1 (30–60cm), level2 (80-100cm), level3 (180-250cm), level4 (300-320cm). b) Female response to unpredictable variation in predation risk - Coefficient of variation in egg densities is higher for medium sized class pools (level3 =180-250cm) than smallest pools (level1 = 30-60cm) at low background predation risk. A similar trend is observed at high background predation risk. No eggs were laid in level4 pools. Low and High represent background predation risk and error bars represent bootstrapped 95% confidence intervals.

The CV in egg densities varied to some extent with pool size – small pools (level 1 pools, <50cm) showed a low mean CV, and medium sized pools (level 3 pools, <120cm) showed a higher mean CV under the background of low predation risk. This trend was weak under high predation risk background. We did not find any detectable differences in CV between predator and predator-free pools (Fig. 3b)

## DISCUSSION

By taking the rare approach of measuring natural variation in predation risk in the wild, our study demonstrates that the wide variation in female OSS behaviour is an outcome of ovipositing females showing a multi-level response, sensitive to both predictable and unpredictable variation in larval predation risk. We demonstrate that females are capable of tailoring responses to predictable variation in predation risk - avoiding large pools that naturally harbour predators throughout the mosquito breeding season; and showing a response sensitive to background predation risk for small pools. Females appeared to respond towards unpredictable variation in larval predation risk by showing a highly variable oviposition response towards medium sized pools that vary stochastically in harbouring predators.

Theory and empirical work suggest that when risk varies predictably in the environment, animals are likely to gain fitness benefits by showing a risk-sensitive response (Heithaus and Dill 2006, reviewed in Verdonlin 2006, Hebblewhite and Merrill 2009). However, most studies on anti-predation responses rarely quantify both predictable and unpredictable natural variation in predation risk over space and time, *a priori*, to test anti-predation responses. Studies typically vary the concentration of predator cues in a controlled experimental set-up and measure immediate behavioural avoidance responses (Juliana, Kotler, Brown, Mukherjee, & Bouskila, 2011; Brown, Rive, Ferrari, & Chivers, 2006; Brown, Ferrari, Elvidge, Ramnarine, & Chivers, 2013). The mosquito model system allowed us to first quantify larval predation risk in space and time in the wild, and then test behavioural responses through manipulative field experiments. By measuring predation risk in the wild, we were able to identify predictable and unpredictable elements of variation in predation risk. We identified clear predictable variation in predation risk – both in space among pools of different sizes and in time as the mosquito breeding season progressed. Predictable sources of variation in predation risk was found to exist in the form of spatial and temporal variation. Spatial variation was between pools distributed in space – small pools reliably predicted low risk and large pools reliably predicted high predation risk. Temporal predictable variation was in the form of seasonal changes in overall predation risk – low predation risk early in the mosquito breeding season (July to September) followed by high predation risk in October and November. These data on predator densities suggests that early in the season, females make oviposition decisions in a background of low predation risk, but later in the season, females must select among pools in a background of high level of predation risk. These systematic patterns in space and time appear to influence female oviposition responses. Females consistently avoided ovipositing in large pools (irrespective of experimentally induced predator presence/absence) throughout the mosquito breeding season, but not in small and intermediate pools. In the natural setting, including in our study, large pools contain predators throughout the mosquito breeding period, as their high water-holding capacity, makes them suitable habitats for predators that typically have longer development times (Chase & Knight, 2003). Since *Aedes spp.* females are known to naturally oviposit in the size range of pools that we sampled in our study (including the large sized pools: Clements, 1999, personal communication by Mukherjee, personal observation by Sharma), we propose that ovipositing females might be using large pool size as a proxy to gauge predator presence reliably and have been selected to avoid ovipositing in them. Similar evidence for prey animals using indirect cues to avoid predators is provided by several studies from rodents where they show that prey commonly use indirect cues, such as microhabitat features, to assess and avoid predator dense sites (Dill, 1987; Lima, 1987; Lima & Dill, 1990).

Temporal variation in risk appeared to influence female oviposition behaviour. Our findings indicated that depending on the level of background predation risk, females changed the way they responded to immediate predator presence/absence in small to medium pools. Using indirect cues, such as habitat features, is a common strategy to avoid predators at a broad scale (Gilliam & Fraser, 1987; Brown & Kotler, 2004; Kotler, Brown, & Hasson, 1991), whereas, animals are also likely to track variation in predation risk at a much finer scale in space and time (Thaker et al., 2011). Controlled experimental studies show that prey response could be shaped by an interplay between immediate and background predation risk (Brown, Chivers, Elvidge, Jackson, & Ferrari, 2014). In our study with a wild population, we see a similar pattern where oviposition behaviour was found be sensitive to both, the level of background predation risk as well as immediate predator presence/absence status at the experimental pool level. Females preferred ovipositing in predator-free pools when the background predation risk was low, possibly because they gain benefits of increased offspring survival by avoiding predator pools. But, this response switched when the background predation risk was high, possibly because most pools in the background contained a high density of predators. When there are predators everywhere why not lay eggs in predator-free pools? We speculate that the search cost of locating a predator-free pool might outweigh the benefits of ovipositing in a predator-free pool. However, this prediction needs to be tested in a controlled experiment by measuring protein and lipid reserves spent on search flight. This conditional response could also be explained by other factors that vary as the season progresses which we could not control in the experiment, such as predation on adult mosquitoes or changes in ovipositing female densities. Our findings, thus, demonstrate that females track variability in predation risk not just at the level of pools, but also at the level of multiple pools by paying attention to the background predation risk represented by the network of pools on the sheet-rock.

An additional source of predictable variation that females appeared to respond to is seasonality in breeding condition relating to diapause season. Females showed a seasonal pattern in oviposition - in the diapause season (when eggs undergo dormancy), females oviposited indiscriminately with respect to predator presence/absence, and laid a much higher number of eggs than in the pre-diapause season. Similar seasonal changes in oviposition behavior is found in many insect groups, where they track photoperiod length, temperature, to respond to seasonal variation in environmental factors (Tauber & Tauber, 1976)), Edgerly, McFarland, & Morgan, 1998). There is a growing body of literature that shows that animals track predictable variation in ecological factors at multiple spatial scales and vary these responses along the season. Animals tracking variation at multiple spatial scales has been shown in different suites of behaviour: habitat selection for breeding (Chalfoun & Martin, 2007), Sharma et al., under review); foraging patch selection varied with abundance of food resources (Naniwadekar, Mishra, & Datta, 2015). Tracking seasonal variation is reported from many studies, where animals are shown to be sensitive to changes in ecological factors along the season, for example, breeding decisions in many birds are influenced by seasonal changes in larval food resources (Eeva, Veistola, & Lehikoinen, 2000), microhabitat use decisions in many aquatic insects are influenced by seasonal changes in temperature and humidity (Refsnider & Janzen, 2010).

How do females tackle unpredictable variation, i.e.,when the risk varies in a stochastic manner such that there is no reliable relationship between the signal (in our case, predation risk) and fitness consequences of animal response? In the first part of our study, we measured variation in predation risk in the wild, and we identified medium sized pools to be associated with unpredictable variation in predation risk. From the manipulative pool experiments, we find some evidence that female response towards medium sized pools, that are inherently stochastic in predicting predation risk, was more variable than their response towards small and large pools. Currently, we do not know much about the manner in which unpredictability influences oviposition decisions. Theory suggests that unpredictability in the environment could be an erosive force for trait evolution, as extreme phenotypes at the tails of the phenotypic distribution would be eliminated at each generation (Lenormand, Roze, & Rousset, 2009), for example, a wet specialist or a dry specialist might not do well in the long term if drought and wet periods occur stochastically in the environment (Starrfelt & Kokko, 2012). One explanation for the variation in oviposition response in our study could that females show random oviposition, i.e., they oviposit in pools that they encounter at random. This could indicate lack of selection through erosion of extreme phenotypes, in our case, complete avoidance of predator pools or preference for predator-free pools. A second explanation is the evolution of alternative bet-hedging strategies for oviposition as an adaptive response towards long-term stochasticity. From controlled experiments in our laboratory using individual females from a colony of *Aedes aegypti*, we find evidence for egg spreading strategy (Chaithra, unpublished data). Other studies have reported similar findings where *Aedes* spp. distribute eggs between oviposition sites (Clements, 1999) and we speculate that this could be a mechanism to spread risk in response to unreliable cues of future predator presence. Many theoretical studies have provided the basis for the idea that in unpredictable environments, natural selection might favour strategies that reduce variance in fitness although at the expense of a reduced mean fitness (Seger & Brockmann, 1987), but empirical evidence for bet-hedging strategies is still very limited. A few studies have alluded to bet-hedging strategies as a mechanism to explain unaccountable variation, such as diversification in timing of seed germination (Philippi, 1993) or occurrence of multiple matings (Byrne & Keogh, 2009), evolution of iteroparity (Cunnington & Brooks, 1996) or high ovule number per flower (Rees, Childs, Rose, & Grubb, 2004). Empirical evidence for bet-hedging is limited (but see Rees, Childs, Rose, & Grubb, 2004) as multi-generational studies are usually required to establish the adaptiveness of putative bet-hedging traits (Simons, 2011, Childs et al. 2010). Our study provides a useful starting point to test evolution under stochastic environments in the future, i.e., to identify unpredictable variation in predation risk in the wild.

Our study also shows that apart from examining variation in a selective factor, we also need to incorporate the effects of other important selective factors. A clear result in our study was that females did not prefer to oviposit in small pools that are naturally devoid of larval predators throughout the season. The hydroperiod length of small pools is too short to support predators and yet, females did not show a clear preference for these pools (<50cm perimeter). This finding was explained well by our results from examining variation in pool desiccation risk. Our measures on pool hydroperiod length suggest that in these small pools the risk of desiccation is high for larvae, as these pools are likely to desiccate before the *Ae. vexans* larvae can complete their maturation phase of 4-5 days (Clements, 1999). Thus, the benefits from avoiding predators may be outweighed by the costs from desiccation risk, suggesting that other selection pressures may interact with predation risk to shape oviposition site selection decisions.

Our study attempts to emphasise that seemingly puzzling variation in a behaviour can be explained once we carefully build a framework for the multiple ecological processes likely to affect that behaviour in natural ecological contexts. Using this framework, we show that female oviposition behaviour is complex - 1) females avoided large pools that permanently harboured predators in natural settings, suggesting they could be using large pool size as an indirect proxy for high risk of larval predation in future, 2) females did not consistently discriminate between pools based on direct predator presence, this response was sensitive to background predation risk, 3) females showed higher variation in oviposition behaviour when faced with naturally stochastic medium sized pools, highlighting the role of unpredictability in predation risk in shaping female behaviour, 4) females did not show a strong preference for small pools that are naturally are devoid of predators suggesting that females balance both predation and desiccation risks when choosing oviposition sites, and 5) female oviposition is sensitive to diapause season. Taken together, these findings show that oviposition site selection in *Aedes vexans*, is likely to be shaped by two processes - multiple selective factors, and variation, both predictable and unpredictable, in the selective factors. Studies rarely make detailed quantitative measurements of variation in environmental factors influencing a trait of interest; consequently, these studies miss on information particularly on stochastically varying environmental factors. We suggest future studies to focus measuring the natural variation in predation risk with an aim to identify unpredictable variation in predation risk to study prey trait evolution.

